## Supplementary material for "Bridging the gap: Using reservoir ecology and human serosurveys to estimate Lassa virus spillover in West Africa": S1 Appendix

#### 1 Feature data

We obtained land cover classification rasters for West Africa from the Moderate Resolution Imaging Spectroradiometer (MODIS) dataset [1]. The *Mastomys natalensis* occurrences, as well as occurrences of Lassa virus (LASV) in *M. natalensis*, were collected between 1972 – 2017. However, MODIS products were not available prior to 2001, and as a result, finding any MODIS predictors that line up with the year in which a rodent capture or Lassa survey occurred was, in general, not possible.

Because it is likely that certain land features described by the MODIS data have changed throughout this time interval, we use MODIS land cover data from 2001, and only include as predictors those land cover types that were least likely to change throughout the time interval over which the response variables were collected. To find which land cover types changed the least, we first downloaded all land cover data for West Africa for 2001 – 2018. Let $L$ denote the set of possible land cover types. For each year between 2001 and 2017, and for every pixel across West Africa, we calculated the frequency of land cover transitions from type $j \in L$, to type $i \in L$. Letting $X_t$ denote the land cover type of an arbitrary pixel during year $t$, these frequencies allow us to estimate a land cover transition matrix

$$M_{i,j} = \Pr(X_{t+1} = i | X_t = j). \tag{1}$$

With the matrix M in hand, we can estimate the probability that a pixel of land cover type $i$ changes over the course of a single year (i.e. $X_t = i \neq X_{t+1}$):

$$C_i = 1 - M_{i,i}. \tag{2}$$

Finally, we use $C_i$ to calculate the average duration of each land cover type. Assuming that, for a given pixel, the times of land cover transitions are geometrically distributed, the duration in years of land cover type $i$ is

$$T_i = \frac{1}{C_i}. \tag{3}$$

From the 2001 MODIS land cover data, we include predictors with an expected duration of at least 20 years: $\{i \in L \,|\, T_i \geq 20\}$. This criterion allows eleven different land cover features in the model: evergreen broadleaf forest, water bodies, open shrubland, woody savanna, savanna, grassland, permanent wetland, cropland, urban builtup, cropland natural mosaic, and barren.

In its original form, the raster data classifies each pixel into discrete land cover types. Because the coordinates of our spatial point data are not known with absolute certainty, we modified the original land cover raster product into a raster stack with eleven layers that describe the local density of each land cover type in the surrounding area. For example, an arbitrary focal pixel of the first layer describes the fraction of 0.05° pixels that, within the surrounding 0.15° x 0.15° area, are classified as "evergreen broadleaf forest".

In addition to land cover data, we also obtained monthly rasters that describe temperature, rainfall, and Normalized Difference Vegetation Index (NDVI) between 2001 – 2019. Temperature and NDVI was obtained from the MODIS source at monthly resolution [2, 3]. Rasters of monthly averaged precipitation were obtained from the Climate Hazards Group InfraRed Precipitation with Station data (CHIRPS) dataset [4].

Rainfall, temperature, and NDVI datasets were used to calculate several measures of environmental seasonality. First, these data were processed to generate three rasters that describe corresponding mean values across all years between 2001 and 2019. Because of missing values in the temperature data, the following derived variables were only calculated for rainfall and NDVI. The coefficient of variation was calculated within each year, and averaged over the 19 years of data. We found the average maximum precipitation that was obtained within any year, as well as the average minimum. These same predictors were calculated for NDVI. We also included measurements of Colwell's indices for precipitation and NDVI, including both constancy and contingency, which together describe the predictability of a seasonal phenomenon [5]. A high precipitation constancy, for example, indicates a region in which monthly rainfall is predictable because it is relatively constant. In contrast, a region with a seasonal wet season has lower constancy of precipitation, but is still predictable because of its periodicity. Such a region has a high contingency [5]. In addition, we included measures of the duration of low precipitation for each year, defined as months with daily precipitation of less than 1 mm/day, and averaged these over the 19 years of data. A similar feature was calculated for NDVI using a threshold value of 0.5.

A raster of elevation in meters of West Africa was obtained from the digital elevation model (DEM) dataset from the USGS (U.S. Geological Survey Center for Earth Resources Observation and Science) [6]. Lastly, we included the projected 2020 population raster of Africa from the WorldPop dataset [7]. This data was used to derive estimates of human density in each 0.05° x 0.05° pixel. We chose population data projected for the year 2020 because, although population densities across West Africa changed throughout the time-frame of the response variables, the relative population densities between locations in West Africa are broadly similar to previous years. Furthermore, because population density plays an important role in predicting the rate of new cases, the most up-to-date population data will provide the best estimates of current spillover risk from Lassa.

#### 2 Predictors used by the fitted models

Fig 1 shows the ten environmental features that best lowered the model deviance across all 25 bootstrap fits of the $D_M$ layer. The top four predictors were NDVI contingency, NDVI coefficient of variation, mean precipitation, and precipitation coefficient of variation. Fig 2 shows the effect of the top six features on the belief that a pixel contains *M. natalensis*. The models learned to positively associate the presence of *M. natalensis* with relatively high values of NDVI contingency, indicating that the species thrives in places that have strong, seasonal patterns of vegetation. In addition, the algorithm assigned greater predictions in locations with particular rainfall patterns, including a specific mean and max range of precipitation.

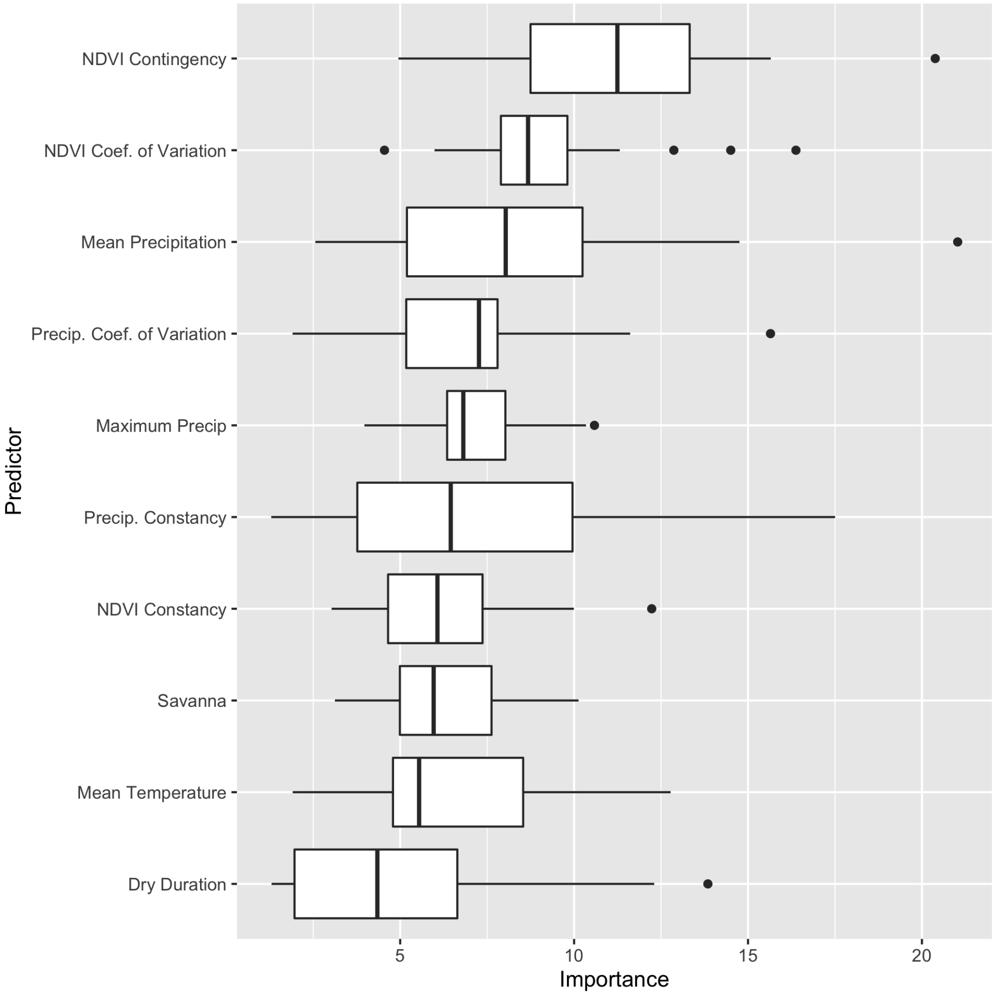


**Fig 1. Influential predictors in the reservoir layer.** Top ten most influential environmental features, chosen across 25 bootstrapped models, that discriminate *M. natalensis* presences from background points. Here, importance measures the relative average extent to which inclusion of a feature variable lowered the estimated deviance across all bootstrap fits.

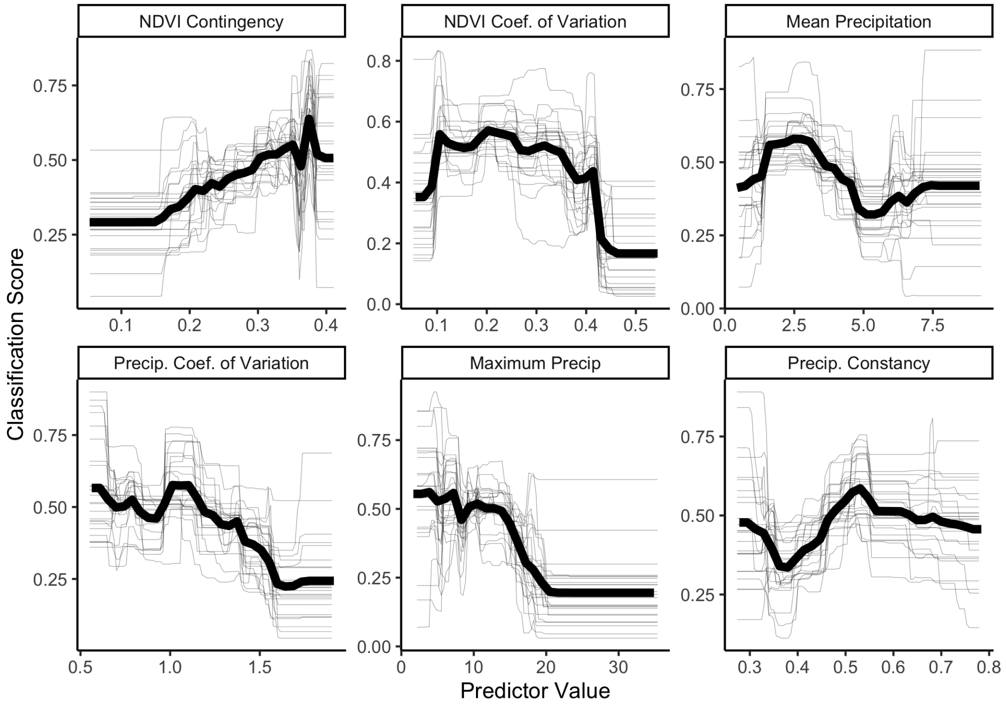


**Fig 2. Environmental relationships learned by the reservoir layer.** Shown for the six predictors that best discriminated *M. natalensis* presences from background points. For a given panel, the x-axis indicates the predictor value, and the y-axis shows the classification score that is assigned (averaged over all other predictor values). Each thin line shows the relationship of one bootstrap simulation; thick black line shows the mean across all 25 models.

Fig 3 shows the top ten features chosen by the $D_L$ layer's tree-fitting algorithm across 25 bootstrap model fits. The algorithm primarily used precipitation contingency to determine whether or not a pixel is suitable for endemic LASV in *M. natalensis*. Fig 4 shows the effect of the top six predictors. The fitted model's belief that a pixel has harbored LASV was positively impacted by areas with high precipitation contingency (i.e., predictable rainfall patterns).

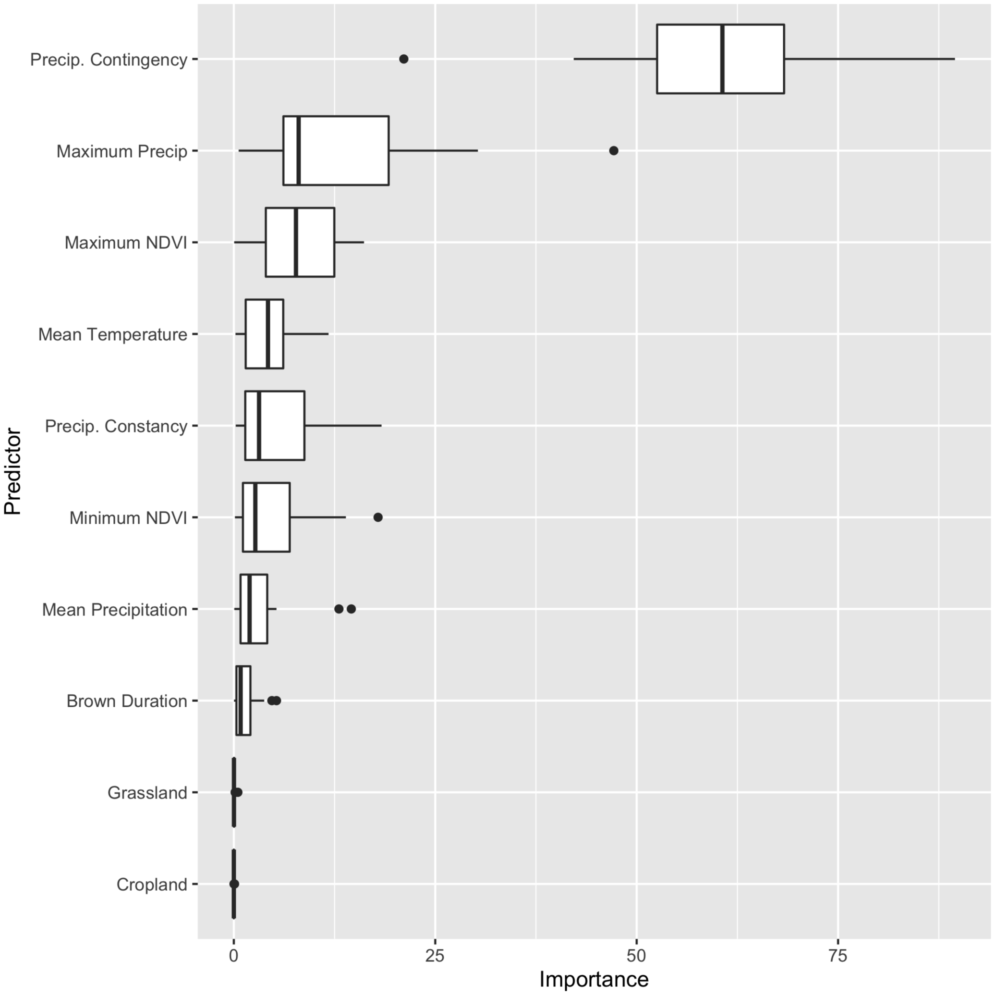


**Fig 3. Top ten most influential features chosen by the pathogen layer.** Chosen across 25 bootstrapped model fits, these are the predictors that discriminate between surveyed pixels with vs without LASV-infected *M. natalensis*. Importance measures the relative average extent to which inclusion of a feature variable lowered the estimated deviance.

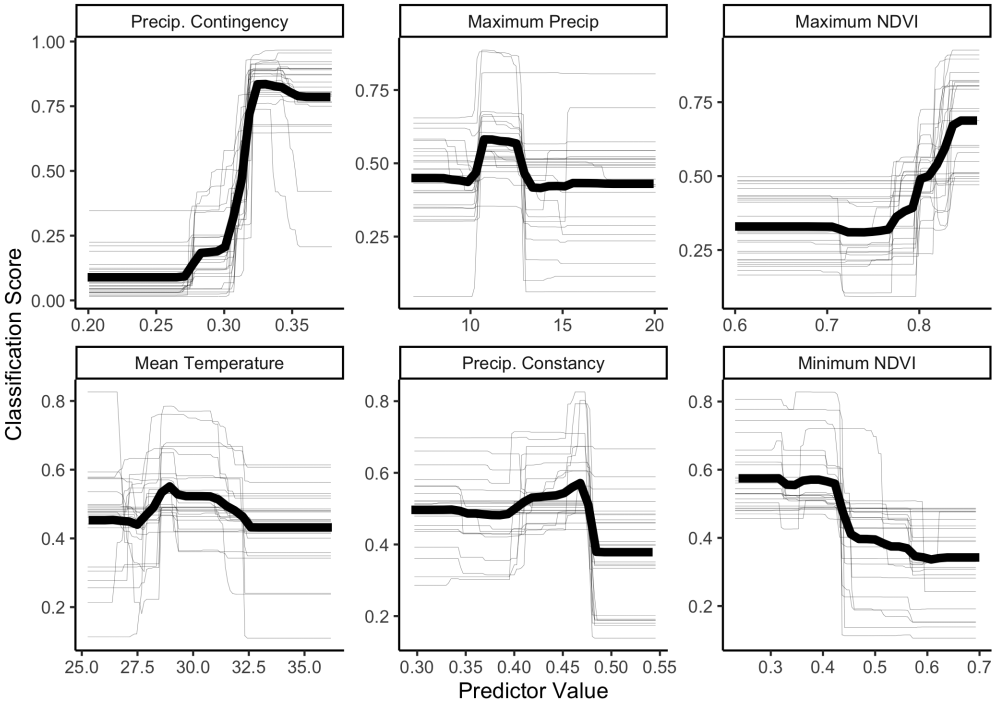


**Fig 4. Environmental relationships learned by the pathogen layer.** Six most influential predictors that best discriminate between presence vs absence of LASV in *M. natalensis*. Each thin line shows the relationship of one bootstrap simulation; thick black line shows the mean across all 25 models.

### 3 Generalized Linear Regression of Human Seroprevalence

Here, we present the statistical output of the quasi-binomial regression of human seroprevalence onto spillover risk $D_X$ and an intercept. As preliminary work, we first fit a binomial regression with each seroprevalence estimate weighted by the number of individuals that were tested. The fitting procedure resulted in a null deviance of 1613.8 (93 degrees of freedom) and a residual deviance of 1361.1 (92 degrees of freedom). A plot of the deviance residuals indicated homoskedastic variability (Table 1, Fig 5). However, because most of the deviance residuals are outside the range $[-1, 1]$, the fitted model also implied that the variability of the residuals is greater than that assumed in a standard binomial regression (i.e., are overdispersed). Overdispersion, in turn, contaminates the hypothesis testing of model coefficients.

**Table 1. Distribution of the deviance residuals in the binomial regression.**

| Min | 1Q | Median | 3Q | Max |
|---|---|---|---|---|
| -9.1 | -2.9 | -1.4 | 2.3 | 8.7 |

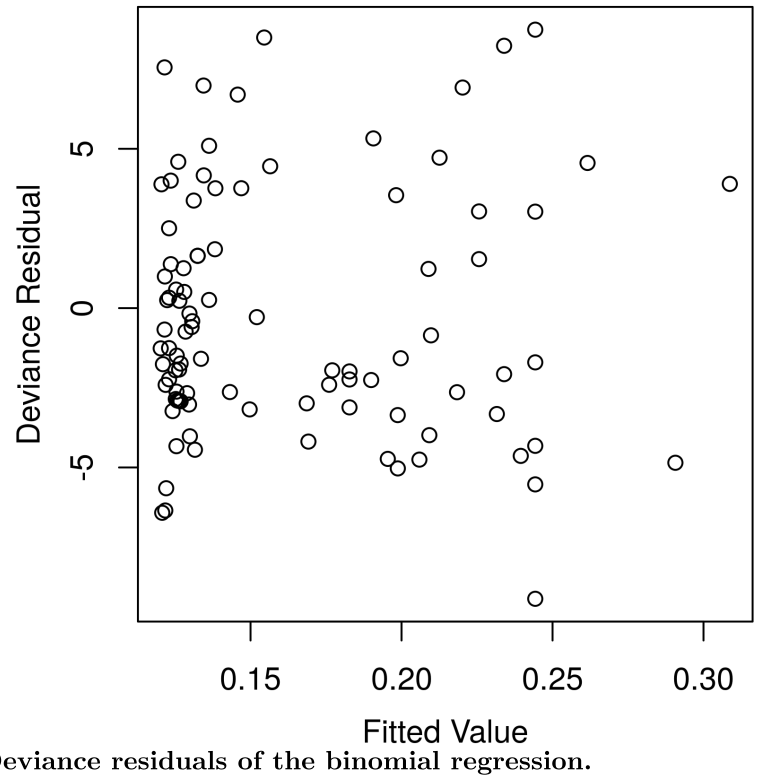


**Fig 5. Deviance residuals of the binomial regression.**

We fit a quasi-binomial regression to account for the overdispersion. As in the binomial regression, each seroprevalence estimate was weighted by the number of individuals tested. The quasi-binomial regression yields the same coefficient estimates and predictions as a binomial regression, but also estimates a dispersion parameter that accounts for increased variance in the residuals. The dispersion parameter was estimated to be 15.1. This estimate is used to correct confidence intervals on the model coefficients by a scaling factor $\sqrt{15.1}$. Using these corrected confidence intervals, the regression indicated that spillover risk ($D_X$) was significantly associated with human seroprevalence (Table 2).

**Table 2. Coefficient estimates and hypothesis tests for the generalized linear regression.**

| | Estimate | Std. Error | t-value | Pr($>|t|$) |
|---|---|---|---|---|
| Intercept | -2.00 | 0.17 | -11.9 | $<$ 2e-16 |
| Dx | 1.50 | 0.37 | 4.01 | 0.000123 |

#### References


1. Friedl M, Sulla-Menashe D. MCD12C1 MODIS/Terra+Aqua Land Cover Type Yearly L3 Global 0.05Deg CMG V006 [Data set]; 2015.

2. Didan K, Huete A. MOD13C2 MODIS/Terra Vegetation Indices Monthly L3 Global 0.05Deg CMG V006 [Data set]; 2015.

3. Wan Z, Hook S, Hulley G. MOD11C3 MODIS/Terra Land Surface Temperature/Emissivity Monthly L3 Global 0.05Deg CMG V006 [Data set]; 2015.

4. Funk C, Peterson P, Landsfeld M, Pedreros D, Verdin J, Shukla S, et al. The climate hazards infrared precipitation with stations—a new environmental record for monitoring extremes. Sci Data. 2015;2(1):150066. doi:10.1038/sdata.2015.66.

5. Colwell RK. Predictability, constancy, and contingency of periodic phenomena. Ecology. 1974;55(5):1148–1153. doi:10.2307/1940366.

6. U S Geological Survey Center for earth resources observation and science. GTOPO30 Global 30 Arc-second Elevation; 2018.

7. School of Geography and Environmental Science, University of Southampton; Department of Geography and Geosciences, University of Louisville; Departement de Geographie, Universite de Namur) and Center for International Earth Science Information Network (CIESIN), Columbia University Global High Resolution Population Denominators Project - Funded by The Bill and Melinda Gates Foundation (OPP1134076). WorldPop; 2020.
